# Cyclic dominance emerges from the evolution of two inter-linked cooperative behaviours in the social amoeba

**DOI:** 10.1101/251553

**Authors:** Shota Shibasaki, Masakazu Shimada

## Abstract

Evolution of cooperation has been one of the most important problems in sociobiology, and many researchers have revealed mechanisms that can facilitate the evolution of cooperation. However, most studies deal only with one cooperative behaviour, even though some organisms perform two or more cooperative behaviours. The social amoeba *Dictyostelium discoideum* performs two cooperative behaviours in starvation: fruiting body formation and macrocyst formation. Here, we constructed a model that couples these two behaviours, and we found that the two behaviours are maintained due to the emergence of cyclic dominance, although cooperation cannot evolve if only either of the two behaviours is performed. The common chemoattractant cyclic AMP is used in both fruiting body formation and macrocyst formation, providing a biological context for this coupling. Cyclic dominance emerges regardless of the existence of mating types or spatial structure in the model. In addition, cooperation can re-emerge in the population even after it goes extinct. These results indicate that the two cooperative behaviours of the social amoeba are maintained due to the common chemical signal that underlies both fruiting body formation and macrocyst formation. We demonstrate the importance of coupling multiple games when the underlying behaviours are associated with one another.

## Introduction

Evolution of cooperation has been a challenging topic in sociobiology. Although it is a widespread behaviour, apparent from human societies [7] to microbial communities [55], cooperation is vulnerable to invasion by defectors. Cooperators pay a cost for helping other individuals, while defectors pay no cost but receive the benefits from cooperators. As a consequence, natural selection should favour defectors [29, 56]. However, many studies have revealed mechanisms that facilitate the evolution of cooperation, including kin discrimination [21, 23, 42], spatial structure that enables cooperators to interact frequently with one another [2, 8, 26, 27], and a by-product which increases the benefit for cooperators (e.g., pleiotropy [10, 25, 51, 52]).

Most studies have explored only one cooperative behaviour at a time. However, some organisms, such as the social amoeba *Dictyostelium discoideum*, perform multiple types of cooperation. It has been known that *D. discoideum* performs two distinct types of cooperative behaviours in starvation. One of these is fruiting body formation, wherein some cells differentiate into spore cells and survive while other cells differentiate into nonviable stalk cells and aid in dispersal of spore cells [41]. The offspring germinate from spore cells after conditions become favourable [22]. The other cooperative behaviour is known as macrocyst formation, wherein some cells become sexually mature and produce offspring after forming diploid zygote cells, but other cells remain vegetative and provide energy to the zygote cells through cannibalism [31]. The haploid progeny eventually appears, but the germination process remains unclear [31].

In fruiting body formation, cells first aggregate by responding to synchronously secreted cyclic AMP (cAMP) [13], and then, the aggregated cells form multicellular slugs and develop into fruiting bodies. When cells from two or more strains form a fruiting body (i.e., forming a chimeric fruiting body), the cell-type ratio has a large effect of the fitness of each strain [43, 36]. If cells from one strain differentiate only into spore cells, this strain can receive a large benefit through the formation of chimeric fruiting bodies because they pay no cost for stalk formation. As differentiation is a key to fruiting body formation, theoretical studies have focused on signalling chemicals that induce stalk formation [32, 50].

In macrocyst formation, about 16% of cells first mature sexually and differentiate into gametes or fusion component (FC) cells [30], whereas other cells remain vegetative. In the heterothallic strains of *D. discoideum*, there are three mating types, and each FC cell fuses with that of a different mating type [3]. After cell fusion, pronuclear fusion occurs [18] and zygote cells are formed. The zygote cells then secrete cAMP [1] to collect the vegetative cells around them for cannibalization, which leads to mature zygote cells in the macrocysts. Macrocyst formation can be considered as a form of cooperation because (i) the division of labours between FC cells (producing offspring thorough macrocyst formation) and vegetative cells (offering energy thorough cannibalization) is similar to the relationship between spore cells and stalk cells in fruiting body formation, and (ii) the vegetative cells can avoid the cannibalism and survive if they ignore cAMP from the macrocyst (i.e., a strain behaves as a defector in macrocyst formation). In contrast to fruiting body formation, however, there are fewer studies on the evolution of macrocyst formation, likely due to difficulties in germinating macrocysts in the laboratory [11]. To our knowledge, there are only two theoretical studies on macrocyst formation; one shows that fluctuations in food availability play an important role in maintaining the ability to aggregate the vegetative cells [6], and the other indicates that without food fluctuation, vegetative cells would still respond to cAMP if macrocyst formation co-occurs with fruiting body formation and if cooperation in fruiting body formation is preserved [39].

Here, we use a modeling approach to demonstrate that the two cooperative behaviours of *D. discoideum*, fruiting body formation and macrocyst formation, can be maintained by coupling the evolution of these two behaviours. First, we considered a very simple situation that ignored mating types and spatial structure. In this model, cyclic dominance, a loop structure where one strategy beats another strategy but this strategy is beaten by still another strategy, emerged under some conditions (see inequality (7) and figure 2), and cooperation in both fruiting body formation and macrocyst formation was maintained. We then introduced variation in mating types and assumed that macrocyst formation occurred only between different mating types. Cyclic dominance again emerged in this case. When the model included spatial structure, such that each colony of the social amoeba interacted only with their neighbours, this result held and cooperation was maintained. Thus, by coupling these cooperative behaviours, cyclic dominance emerges and both behaviours are maintained even if it is assumed that neither cooperative behaviour can evolve alone. These results are consistent regardless of mating types and spatial structure introduced to the model if the environment where fruiting body formation co-occurs with macrocysts is assumed.

## Models

In this section, we describe the methods for the simulations we performed. The materials and methods for the experiments conducted in this study can be found in the Supplementary information (SI).

### Model assumptions

As both fruiting bodies and macrocysts are formed under dark and dry conditions [39] (see also our experimental results in SI and figure S9), we assume that *D. discoideum* can perform both fruiting body formation and macrocyst formation. Note that we ignore homologous recombination during macrocyst formation. For simplicity, we first assume that both fruiting body formation and macrocyst formation are represented as a prisoner’s dilemma (PD) game, where the defector is the evolutionarily stable strategy [28]. Although the PD game can be inadequate for conceptualizing fruiting body formation nor macrocyst formation and cooperation can evolve in each game without coupling them in nature, we conceptualize the two games by the PD games to assume the most difficult conditions for cooperators to evolve in each game. We shall show that cooperation can be maintained by coupling the two games under this assumption.

The payoff matrices for the two games are given by

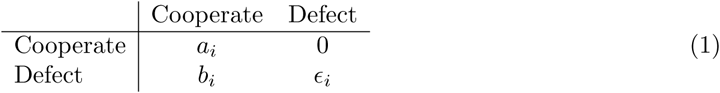

where *i* = 1 corresponds to fruiting body formation, and *i* = 2 refers to macrocyst formation. It should be noted that the inequality *b_i_* > *a_i_* ≫ *ϵ_i_* > 0 is satisfied for *i* = 1, 2 and that cooperation cannot evolve if only either of the two games is performed.

Cooperators of fruiting body formation are those that produce both spore cells and stalk cells, whereas defectors produce only spore cells. When cooperators form fruiting bodies with another cooperator, they receive benefit *a*_1_. On the other hand, defectors receive *b*_1_ when they exploit the cooperators by forming chimeric fruiting bodies, but the cooperator receives no benefit in this case. However, when defectors interact with another defector, they receive only a small benefit *ϵ*_1_ because they both are deficient in stalk formation. Such a mutant strain is referred to as *fbxA*^−^ in *D. discoideum* [12].

In macrocyst formation, cooperators provide energy to zygote cells by making vegetative cells respond to cAMP to allow cannibalization by zygote cells. On the other hand, defectors avoid cannibalistic attack by ignoring cAMP. Such a mutant strain is known as TMC1 [40]. When the two cooperators produce macrocysts together, they each receive a benefit *a*_2_, as macrocysts obtain enough energy through cannibalism. On the other hand, if a defector interacts with a cooperator, the defector receives *b*_2_ by avoiding a cannibalistic attack, but the cooperator receives no benefit. However, when the two defectors form macrocysts together, they receive a very small benefit *ϵ*_2_, as macrocysts cannot receive enough energy from the vegetative cells alone.

It should be noted that the common chemical signal cAMP plays an important role in both fruiting body formation and macrocyst formation; aggregation requires cAMP in both cases. If cells do not aggregate in macrocyst formation (i.e., a defector of macrocyst formation), they do not aggregate in fruiting body formation either. In other words, defectors during macrocyst formation cannot join in fruiting body formation. This phenomenon was experimentally identified [40], and we reflect it in the models (*ϵ*_0_ in payoff matrix given by equation (2)).

We began by building a very simple model, where mating types and spatial structure were ignored. In this case, there were only three strategies: a cooperator of both fruiting body formation and macrocyst formation (*C*), a defector of fruiting body formation that behaved as a cooperator of macrocyst formation (*D^F^*), and a defector of macrocyst formation that could not join in fruiting body formation (*D^M^*). The payoff matrix *A* is given by

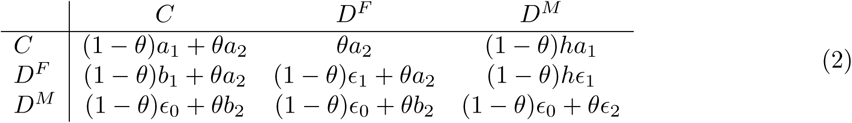

where *θ* is the probability that macrocyst formation occurs, while 1 – *θ* refers to fruiting body formation (0 ≤ *θ* ≤ 1). *ϵ*_0_ is the payoff for remaining as solitary vegetative cells in fruiting body formation. For simplicity, we assume that starvation continues for a long period, where almost all vegetative cells die [9]. In this case, it is reasonable to assume that *ϵ*_0_ ≪ min{*ϵ*_1_,*ϵ*_2_}. *h* denotes the payoff effect for forming small fruiting bodies (0 < *h* ≤ 1). When a cooperator of both fruiting body formation (*C*) (or a defector of fruiting body formation (*D^F^*)) forms fruiting bodies with a defector of macrocyst formation (*D^M^*), *D^M^* cannot join in fruiting body formation because it cannot respond to cAMP. The number of cells within fruiting bodies is thus smaller than that when *C* or *D^F^* interacts with either of them. Such fruiting bodies would be shorter and provide lower payoffs due to the inefficiency of dispersal. The roles for additional parameters *θ, ϵ*_0_, and *h* are shown in figure 1.

**Figure 1:**
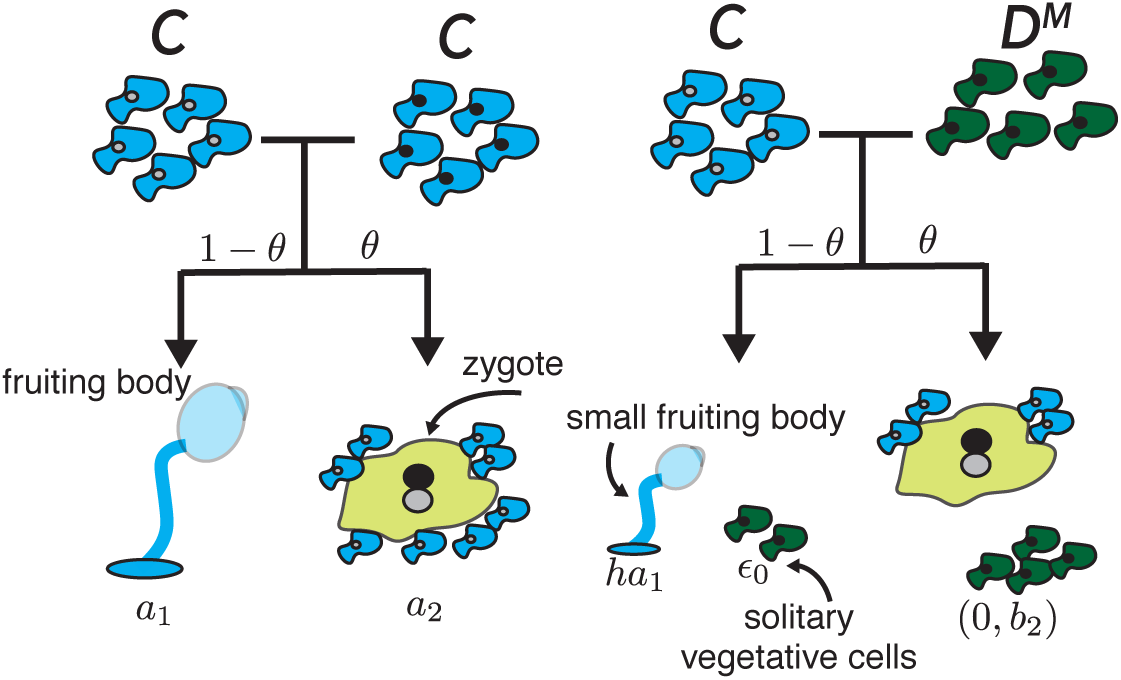
Summary of the additional parameters *θ, ϵ*_0_, and *h*. When the two strategies interact, they form macrocyst formation with a probability of *θ* or fruiting body formation with a probability of 1 – *θ*. If *C* forms chimeric fruiting bodies with *D^M^, C* receives the payoff *ha*_1_ because fruiting bodies are shorter than when two *C*s form fruiting bodies (compare the left case), while *D^M^* receives *ϵ*_0_, as it cannot join in fruiting body formation. Note that *D^M^* can exploit *C* in macrocyst formation, and *D^M^* receives *b*_2_, while *C* receives no benefits in this case.

Due to the constraint of the responsibility to cAMP, the game of fruiting body formation is no longer a PD game. It would be better to call the third strategy *D^M^* as a solitary or a loner in the game of fruiting body formation, which never joins in the game and always receives the benefit *ϵ*_0_ regardless of the strategy of the counterpart. Even in this extended PD game, however, the defect *D^F^* is still the evolutionarily stable strategy, which can be derived by equation (2) when *θ* = 0.

Next, we introduced mating types (type 1 and type 2). In this model, there exist six strategies (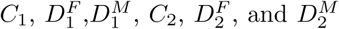 and 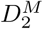), where the subscripts indicate the mating types of each strategy (e.g., *C*_1_ represents a cooperator with mating type 1). Within the mating types, only fruiting body formation is performed, whereas both fruiting body formation and macrocyst formation are performed between the mating types. Note that *θ* represents the probability that macrocyst formation is performed between the mating types. In the model including mating types, therefore, the payoff matrix *S* is described as

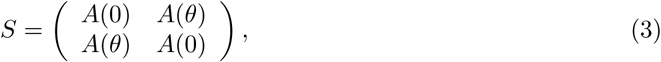

where *A*(0) is a 3 × 3 payoff matrix for the same mating type strategies (they perform only fruiting body formation), while *A*(*θ*) refers to the payoff matrix for different mating type players that perform macrocyst formation with the probability of *θ*.

It should be noted that we ignore the failure of macrocyst formation: i.e., producing FC cells when interacting with the same mating type strains. This assumption will not change the main results in this paper because even if a strain produces FC cell in error, the strain can obtain the benefit from fruiting body formation. In macrocyst formation, some cells differentiate into FC cell while the other cells remain vegetative cells. Such vegetative cells should be able to perform fruiting body formation. Therefore, the strain can receive the benefit from fruiting body formation in the case of failure in macrocyst formation, although the benefits should be smaller than *a*_1_ due to the cost of producing FC cells, which will die at the end. This cost should be small because the ratio of FC cells is about about 16% [30]. Therefore, the failure of macrocyst formation will not change our main results below.

### Replicator Dynamics

The continuous and discrete replicator dynamics are defined, respectively:

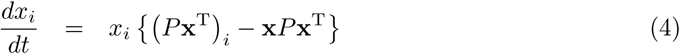

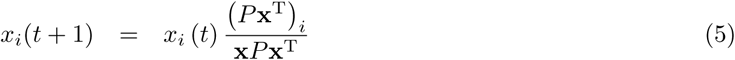

where *x_i_* is the frequency of strategy *i*, x is a vector of the frequencies of each strategy (x = (*x_i_*)), and *P* is a payoff matrix. When the mating types are ignored, *P* equals to the payoff matrix *A* given by equation (2) while *P* equals to the payoff matrix *S* by equation (3) if mating types are included. Numerical computation for continuous replicator dynamics was conducted in Python using the integrators of “ode” with “dopri5” or “odeint” in the Scipy package [19].

### Agent-Based Model

One of the advantages of using an agent-based model is to introduce spatial structure. As macrocyst formation occurs only between mating types, spatial structure can have an effect on the frequencies of fruiting body formation and macrocyst formation. For example, only fruiting body formation would occur if all colonies in the neighbourhood have the same mating type. In addition, the spatial structure would be important for considering the social amoeba in nature, because they perform both fruiting body formation and macrocyst formation though aggregation. In other words, the cells interact only with their neighbours. From these two reasons, we analysed the model with the spatial structure.

In the agent-based model, each agent represents a small colony of *D. discoideum* that has one of the six strategies (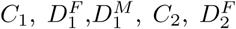 and 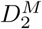) because mating types are also included in this model. By introducing the torus lattice model and Moore neighbourhood, each agent interacts with their eight neighbours at one time step. After all agents finish their interaction and their total payoffs are calculated, they synchronously update their strategies to those of their neighbours with some probabilities given below. In many studies, the Fermi function *f* (*w*) = 1/ {1 + exp(–*w*)} is used as a stochastic update process [44, 53], but the Fermi function is for the pairwise update process. Therefore, we defined the probability Π*_ij_* that agent *i* adopts the strategy of agent *j* using the softmax function

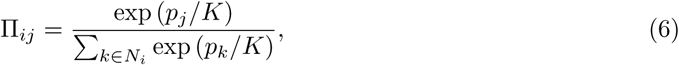

where *p_i_* is the total payoff of agent *i, N_i_* is the set of agents in the neighbourhood of agent *i* (including agent *i* itself), and *K* is the inverse of the intensity of selection. It is more likely that agent *i* adopts the strategy of agent *j* in the next time step if *p_i_* < *p_j_*, but agent *i* stochastically may adopt the strategy of agent *l* even if *p_i_* > *pl*. At the *K* → 0 limit, agent *i* adopts only the strategy of the agent whose payoff is the highest in the neighbourhood. By contrast, as *K* increases, it is more likely that each agent randomly adopts a strategy in their neighbourhood (selection does not work in the *K* → ∞ limit).

We performed the pairwise invasion analysis at 100 × 100 lattice points. One of the two strategies is called an invader, whose initial frequency is 0.01, and the other is a resident. We ran the simulation for 1,000 time steps at various parameter values, and we replicated the simulation 50 times in each case. We then investigated whether the invaders excluded the residents (the invader frequency was greater than 0.99 at the end of the simulation) or they were maintained (the frequencies were greater than 0.01). If invaders excluded the residents, this indicated that the invaders beat the residents. If the invaders did not exclude the residents but they were maintained, they coexisted with the residents.

We then simulated the situations where all six strategies existed at the initial conditions. In this case, we used 300 × 300 lattice points to reduce stochastic effects. In each simulation, the initial frequencies of all strategies were equal, and simulations were continued for 100, 000 time steps. We replicated the simulation 10 times at each value of *θ*.

We also investigated the effect of mutation in the agent-based model. We again used 300 × 300 lattice points, but here we assumed that mutation was rare; at each 500 time step, one agent was randomly chosen to update the strategy to one of the other five strategies, which was again randomly chosen. All codes for the agent-based model were written in C language.

### Data Availability

The simulation codes, the Python script for the analysis, and the experimental data are available at GitHub (https://github.com/ShotaSHIBASAKI/Cyclic-dominance-emerges).

## Results

### Excluding mating types and spatial structure

First of all, we built the simple model that does not include mating types nor spatial structure. In this case, payoff matrix given by equation (2) shows a rock-scissors-paper (RSP) game, approximately if

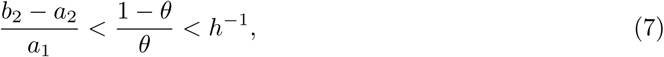

see SI for derivation. In other words, when inequality (7) holds, cyclic dominance emerges, where *D^F^* beats *C, D^M^* beats *D^F^*, and *C* beats *D^M^*. This result can be explained intuitively as follows: in fruiting body formation, *C* is exploited by *D^F^* because fruiting body formation is assumed to be represented as the extended PD game (equation (2) when *θ* = 0). By contrast, in macrocyst formation, *D^F^* is exploited by *D^M^* because we assume that macrocyst formation can be represented by the PD game (equation (1)). It is true that *C* is also exploited by *D^M^* in macrocyst formation, but *C* can enjoy fruiting body formation without exploitation by *D^M^*, which cannot aggregate in fruiting body formation. Therefore, *C* can beat *D^M^* if the payoff of forming fruiting body and the probability of fruiting body formation are sufficiently large. However, if the probability of fruiting body formation is too large, *D^M^* cannot beat *D^F^* as *D^M^* does not obtain a large benefit.

In addition, we found that the continuous replicator dynamics [48] has a unique internal equilibrium *G*, to which any other interior point converges if inequality (7) holds (figure 2, see SI for detail). In this case, a cooperator of both fruiting body formation and macrocyst formation *C* is maintained, although it is not an evolutionarily stable strategy [24]. Therefore, cooperation is maintained if there exist two games, even though cooperation cannot evolve when the social amoeba plays ether of the two games. Other examples of the evolutionary dynamics in this model are shown in figure S1.

**Figure 2:**
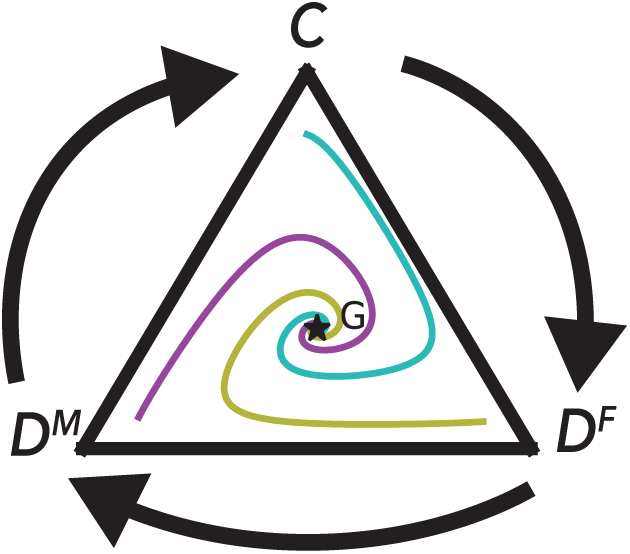
The evolutionary dynamics given by continuous replicator dynamics with payoff matrix by equation (2) when inequality (7) holds. The three orbits (cyan, yellow, and magenta) begin at different initial points but converge at the internal equilibrium *G*. The parameters are *a*_1_ = 1, *b*_1_ = 1.5, *ϵ*_1_ = 10^−4^, *a*_2_ = 0.5, *b*_2_ = 1.2, *ϵ*_2_ = 10^−4^, *θ* = 0.5, *h* = 0.5, and *ϵ*_0_ = 10^−6^.

It should be noted, however, that the application of continuous replicator dynamics is unlikely to be valid for *D. discoideum* because fruiting body formation and macrocyst formation occur in discrete generations; fruiting body formation and/or macrocyst formation occur through starvation, with the offspring subsequently emerging. Thus, discrete replicator dynamics should be applied. Mathematically, the continuous replicator dynamics is a limited case of discrete replicator dynamics [5], which means the result shown in figure 2 is not always valid when the discrete replicator dynamics is applied. Indeed, we found that with discrete replicator dynamics, the range wherein any orbit converges to the internal point *G* is narrower than that in the case of continuous replicator dynamics (inequality (7), see also SI and figure S2).

### A model that includes mating types

When the model includes the existence of mating types, cyclic dominance emerges again (figure 3). In this case, fruiting body formation occurs both within and between the mating types, while macrocyst formation occurs only between the mating types. This means that the parameter *θ* is the probability that macrocyst formation occurs between mating types. As shown in figure 3, *D^F^*s beat both Cs and *D^M^*s within the mating types because only fruiting body formation occurs and *D^F^* is the evolutionarily stable strategy in the game of fruiting body formation (equation (2) when *θ* = 0). However, 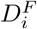 is beaten by the other mating type 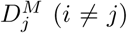 because *D^F^*s are exploited during macrocyst formation and 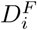 s receive only a small payoff in fruiting body formation, which is not enough for the compensation for macrocyst formation. By contrast, *D^M^*s are beaten by the same mating type Cs because *D^M^*s cannot undergo fruiting body formation due to the lack of the responsibility to cAMP. In addition, under some parameter values, Cs can beat the other mating type *D^F^*s and *D^M^*s (see SI).

**Figure 3:**
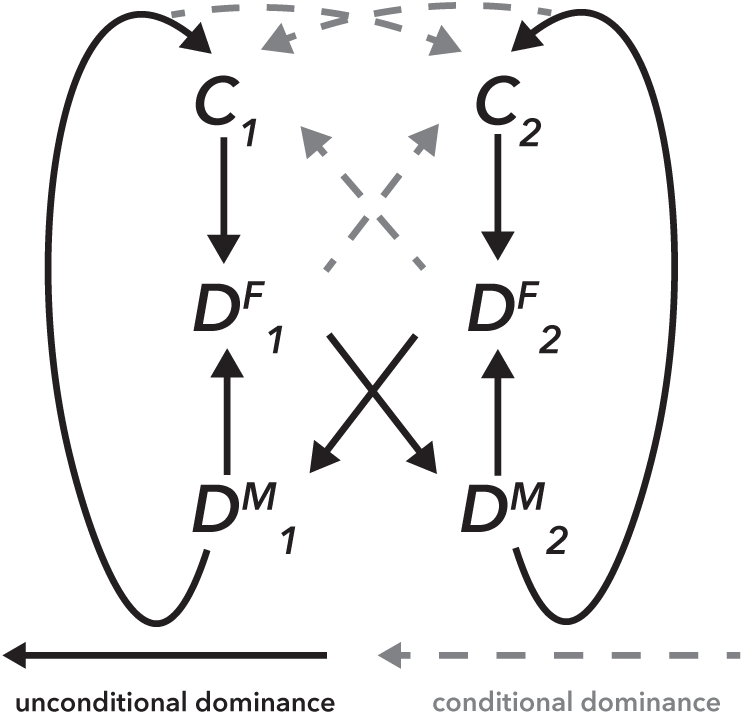
Cyclic dominance emerges when different mating types are introduced. Black arrows show unconditional dominance, and gray arrows show conditional dominance.

Although continuous or discrete replicator dynamics can be applied when mating types are introduced, it is difficult to calculate the stability of an interior equilibrium due to the high dimensionality of the model. In addition, the initial mating type ratio has a large effect on the evolutionary dynamics (figure S3). For these reasons, we used an agent-based model for additional analyses.

### A model that includes spatial structure

In the agent-based model, we again found cyclic dominance that was similar to figure 3, and this result is confirmed by the results of the pairwise invasion analysis shown in figure S4–S6. We then analysed cases wherein all six strategies coexisted equally in the initial conditions. Due to the emergence of cyclic dominance, the cooperators (both *C1* and *C2*) were maintained if *θ* was sufficiently large (figure 4a). When *θ* = 0, only fruiting body formation occurred within and between mating types. As a consequence, defectors of fruiting body formation (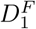 and 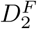) beat the other strategies. If *θ* was small but non-zero (0.1 ≤ *θ* ≤ 0.3), only the defectors of macrocyst formation (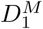 and 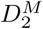) were maintained. This is because the probability of macrocyst formation between mating types *θ* is not sufficiently large for cooperators *C_i_* to obtain the benefits from forming macrocysts with the other mating type cooperators *C_j_* or defectors of fruiting body formation 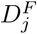 (*j* ≠ *i*) (figures S4), and because defectors of macrocyst formation can exploit other mating type defectors of fruiting body formation (figure S6). If *θ* is sufficiently large (0.4 ≤ *θ* ≤ 1), however, the cooperators can beat the other mating type defectors of fruiting body formation (figure S4) and all six strategies can be maintained (figure 4b). It should be noted that the model with spatial structure do not show the complex dynamics in contrast to the model without spatial structure (figure S3), although the dynamics in the model with the spatial structure shows perturbation.

**Figure 4:**
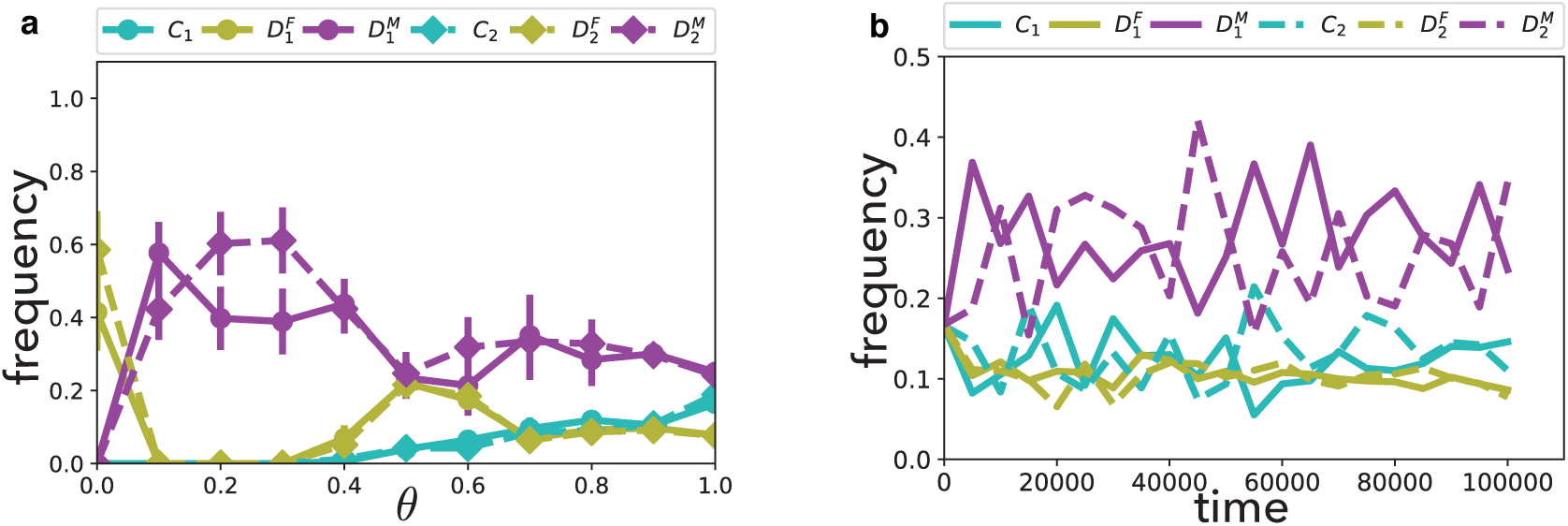
Evolutionary dynamics in the agent-based model. The mean frequencies of each strategy at the end of the simulations with different values of *θ* are shown in (a). The error bars denote standard errors, but some are too small to see. An example of the evolutionary dynamics when *θ* = 0.9 is shown in (b). The parameters are *a*_1_ = 1, *b*_1_ = 2.7, *ϵ*_1_ = 10^−5^, *a*_2_ = 0.5, *b*_2_ = 1.2, *ϵ*_2_ = 10^−5^, *h* = 0.5, *ϵ*_0_ = 10^−6^, and *K* = 1.

In the model with spatial structure, cyclic dominance again contributes the maintenance of cooperation. Although the spatial structure has the positive effect on the evolution of cooperation, we used the parameter values with which Cs goes to extinct if the same mating type *D^F^*s invade (figures S4). In addition, the previous study shows that if the game in macrocyst formation is ignored and cyclic dominance disappears, the coexistence of the two mating types of *D^F^* is the evolutionarily stable state [38]. From these two points, cyclic dominance can be considered to contributing the evolution of cooperation in this model.

Next, mutation was introduced into the agent-based model. Under the assumption that mutation occurs in each 500 time step, only a few strategies can coexist. Here, cooperators can re-invade the population even after they are excluded (figure S8). In the initial condition, two mating types of cooperators (*C*_1_ and *C*_2_) coexist, but these cooperators disappear due to the invasion of the two mating types of *D^F^*s. The coexistence of *D^F^*s is an evolutionarily stable state if the game for macrocyst formation is ignored [38], but this coexistence collapses with the invasion of one mating type defector of macrocyst formation. After that, the either of the two mating types of cooperator appears again by mutation and becomes dominant in the population. Thus, cooperators of both fruiting body formation and macrocyst formation can be revived even after they disappear.

## Discussion

In this paper, our models demonstrate that the two cooperative behaviours of *D. discoideum*, fruiting body formation and macrocyst formation, can be maintained by coupling the two games. While previous research assumed the maintenance of cooperation in fruiting body formation [39], we show that defectors can appear both in fruiting body formation and macrocyst formation. Even though we assume that the defectors are the evolutionary stable strategies in in both of the two games in the single game dynamics, the two cooperative behaviours can be maintained if the model includes both of the games simultaneously.

The primary biological explanation for this result is that the two games are not independent; the common chemical signal cAMP is necessary for aggregation in both fruiting body formation (reviewed in [49]) and macrocyst formation [1]. If some cells do not aggregate (i.e., defect) in macrocyst formation, they receive benefits by avoiding the cannibalistic attack by zygote cells. On the other hand, these cells have substantial costs because they cannot participate in fruiting body formation. Indeed, a mutant that weakly expresses a cAMP receptor will result in poor aggregation for both fruiting body formation and macrocyst formation [40]. This phenomenon is represented as *ϵ*_0_ in the payoff matrix (equation 2), and this provides an explanation why the cooperator of both fruiting body formation and macrocyst formation (*C*) can beat the defector of macrocyst formation (*D^M^*) in the most simple model, wherein the existence of mating types and spatial structure is ignored (figure 2).

The key factor for the maintenance of the cooperators is the emergence of cyclic dominance. Under a simple model without the inclusion of mating types and spatial structure, if the inequality (7) is satisfied, the payoff matrix given by equation (2) follows an RSP game, which is a famous example of cyclic dominance [45]. In addition, the continuous replicator dynamics show the existence of the global attractor in this model (figure 2), and the discrete replicator dynamics shows a similar result (figure S2). Moreover, even when mating types are introduced, cyclic dominance continues to emerge (figure 3). Within mating types, the defectors of fruiting body formation (*D^F^*s) beat the other two strategies because they perform only fruiting body formation. However, *D^F^*s lose to other mating types of the defectors of macrocyst formation (*D^M^*s). Furthermore, *D^M^*s are beaten by the same mating type cooperators *C*, because Cs can undergo fruiting body formation without exploitation by *D^M^*s. The results of the model with spatial structure are also consistent. Cyclic dominance appears again, although all instances are dependent on the parameter values (i.e., conditional advantage, figure S4–S6). Due to cyclic dominance, all six strategies can coexist (figure 4), and cooperators can re-emerge even after they are excluded (figure S8).

The approach we employed in this paper is referred to as multi-games [15, 53, 46] or mixed games [54] in the evolutionary game theory, wherein players play two or more games with different payoff matrices. Although these terms seems unlikely to be used in other fields, it has been known that microbes have multiple inter-linked games or social traits [35]. For example, the production of many kinds of public good is regulated by the quorum-sensing (QS) system [55]. Not only the production of public good, but also the production of QS molecules can be regarded as a game due to the cost of producing QS molecules and the effect of QS molecules other than regulating the production of public goods [33, 37, 57]. In other words, both the production of QS molecules and that of public good are games, and the game of QS molecules has an effect of the public good game.

In addition, recent studies have suggested that microorganisms show cyclic dominance due to the existence of two games. Inglis et al [17] analysed the public good games in *Pseudomonas* communities, wherein there exist two types of public good (i.e., two public good games) and either of the two types of public good is available to each strain. In such communities, the cooperators or the producers can beat the defectors in the other public good game because the defectors cannot exploit the other type of public good, and therefore, cyclic dominance emerges. Another example is shown by Kelsic et al [20], where the the production of antibiotics and their degraders are combined. Although the game of antibiotic production alone can show cyclic dominance, it is impossible to lead the coexistence of all strategies without the spatial structure in the case of the antibiotic production alone. On the other hands, the introduction of the game of degrader production maintains the cyclic dominance and all strategies can coexist without the spatial structure. Considering these studies and our results, multi-game dynamics might be natural in microbial communities and be a factor of stabilizing cooperation and genetic diversity.

It should be noted that our results are based on the assumption that fruiting bodies and macrocysts coexist. Although this has been shown under dark and relatively dry conditions [39], macrocysts are not formed under light conditions [14] and fruiting body formation is inhibited in water [4]. Under these conditions, *D. discoideum* undergoes only one or the other behaviour, and cooperative behaviours are not maintained. In other words, the probability of macrocyst formation *θ* changes over time under changing environments, and cooperation between fruiting body formation and macrocyst formation would be destabilized. Indeed, our model shows that cooperation of fruiting body formation and macrocyst formation is not always maintained under dynamic environments (SI and figure S10). Future research should consider how changing environmental conditions would affect these conclusions.

In addition, our models are a simplification of the complex behaviours exhibited by *D. discoideum*. For example, we assumed distinct strategies (to cooperate or to defect) for both fruiting body formation and macrocyst formation, but these strategies are continuous. Productivity and sensitivity to signals for differentiation in fruiting body formation [32, 50], or the ratio of FC cells and vegetative cells in macrocyst formation [39] can be considered. In addition, we assumed that starvation continues for an extended period of time. If this assumption was relaxed, our results might change because a lack of aggregating cells can be beneficial for fruiting body formation and macrocyst formation if the resource condition quickly recovers [47]. Under this scenario, it may be possible that defectors of macrocyst formation (*D^M^*s) can beat cooperators (*C*s), which would lead to the collapse of cyclic dominance.

Moreover, our results do not contradict with those of previous studies. For example, kin discrimination by the cell adhesion proteins TgrB1 and TgrC1 [16] and the greenbeard effect encoded by *csA* [34] are important for maintenance of cooperation in fruiting body formation. If these effects are included in a model, cooperation would be maintained. Our results, on the other hand, suggest that cooperation can be maintained even if such effects do not exist or if they are incomplete. In addition, if kin discrimination and/or the greenbeard effect are introduced in our models and *D^F^*s do not spread, defectors of macrocyst formation (*D^M^*s) may be unable to invade and only cooperators (*C*s) will be maintained because *D^M^*s cannot beat Cs.

## Conclusion

In summary, we propose that cooperative behaviours in the social amoeba can be maintained by coupling the two games. This is because cyclic dominance emerges from the common cAMP signal that is necessary for aggregation in both behaviours. This result is consistent when mating types and spatial structure are introduced into the model, suggesting the generality of these findings and the importance of including multi-games in behavioural models, especially if the games are inter-linked. Such multi-game dynamics of inter-linked games seems general in microorganisms.

## Authors’ contributions

S.S. and M.S. designed the research, S.S. performed the research and analysed the data, and S.S. and M.S. wrote the paper.

## Acknowledgements

We would thank two anonymous referees for their fruitful comments on the manuscript of the earlier version. We would also like to acknowledge National BioResource Project by the Ministry of Health Labour and Welfare Japan for providing the two strains of *D. discoideum* and *Escherichia coli* b/r.

